# Impact of mutation rate and selection at linked sites on DNA variation across the genomes of humans and other homininae

**DOI:** 10.1101/452201

**Authors:** David Castellano, Adam Eyre-Walker, Kasper Munch

**Author notes:** Centre for Genomic Regulation (CRG), The Barcelona Institute of Science and Technology, Dr. Aiguader 88, Barcelona, 08003, Spain.

## Abstract

DNA diversity varies across the genome of many species. Variation in diversity across a genome might arise from regional variation in the mutation rate, variation in the intensity and mode of natural selection, and regional variation in the recombination rate. We show that both non-coding and non-synonymous diversity are positively correlated to a measure of the mutation rate and the recombination rate and negatively correlated to the density of conserved sequences in 50KB windows across the genomes of humans and non-human homininae. Interestingly, we find that while non-coding diversity is equally affected by these three genomic variables, non-synonymous diversity is mostly dominated by the density of conserved sequences. The positive correlation between diversity and our measure of the mutation rate seems to be largely a direct consequence of regions with higher mutation rates having more diversity. However, the positive correlation with recombination rate and the negative correlation with the density of conserved sequences suggests that selection at linked sites also affect levels of diversity. This is supported by the observation that the ratio of the number of non-synonymous to non-coding polymorphisms is negatively correlated to a measure of the effective population size across the genome. We show these patterns persist even when we restrict our analysis to GC-conservative mutations, demonstrating that the patterns are not driven by GC biased gene conversion. In conclusion, our comparative analyses describe how recombination rate, gene density, and mutation rate interact to produce the patterns of DNA diversity that we observe along the homininae genomes.

## Introduction

The level of genetic variation is known to vary across the genome of many species and this depends on genomic characteristics such as recombination, gene density and mutation rate. This was first demonstrated by Begun and Aquadro (Begun & Aquadro 1992) who showed that putatively neutral genetic diversity was correlated to the rate of recombination across the genome of *Drosophila melanogaster.* This has subsequently been observed in species as diverse as humans and tomatoes (reviewed by (Cutter & Payseur 2013)).

Variation in diversity across the genome of a given species might arise from variation in the mutation rate, selection, and recombination rate. The mutation rate can affect the level of diversity both directly and indirectly. Directly, the level of genetic diversity is expected to depend upon the rate of mutational input; the higher the mutation rate, the more diversity there is expected to be. It can also have an indirect effect by increasing the frequency of selection at linked sites, which is described below. Natural selection can also affect the level of genetic diversity both directly and indirectly. Direct selection tends to either decrease or increase diversity at the sites at which it is acting, depending on whether the selection is either negative or positive, particularly if there is balancing selection. However, in general, selection tends to act indirectly reducing diversity at linked sites through the processes of genetic hitch-hiking (HH) (Smith & Haigh 1974) and background selection (BGS) (Charlesworth et al. 1993). Genetic hitch-hiking also has the effect of moving a locus away from a state of quasi-equilibrium; after a selective sweep, deleterious genetic variation approaches its equilibrium value more rapidly than neutral variation, leading to a disproportionate amount of the diversity being deleterious (Gordo & Dionisio 2005; Pennings et al. 2014; Brandvain & Wright 2016; Do et al. 2015; Castellano et al. 2018). Finally, recombination also affects the level of genetic diversity both directly and indirectly. In humans and all organisms that have been studied, gene conversion appears to be GC-biased (gBGC) (Duret & Galtier 2009; Marais et al. 2001; Pessia et al. 2012; Eyre-Walker 1993; Galtier et al. 2001; Marais 2003). This is counter to the pattern of mutation, which is AT-biased. In such a system in which gBGC and mutation are in opposite direction, an increase in gBGC (from no gBGC) is expected to slightly increase the levels of diversity before reducing them, when gBGC becomes strong (McVean & Charlesworth 1999). However, cross overs induced by recombination tend to act indirectly by reducing the effect of linked selection across loci increasing the local levels of genetic diversity (Berry & Barbadilla 2000).

If selection at linked sites is pervasive and acts genome-wide, this should be visible as correlations between DNA diversity and factors affecting the intensity of selection at linked sites, such as recombination rate, gene density and mutation rate. In this work we return to the question of whether selection at linked sites and mutation rate variation has an effect on levels of DNA sequence diversity and the efficiency of purifying selection along the autosomes of humans and our closest living relatives, the homininae subfamily: humans, bonobos, chimpanzees and gorillas. The role that mutation, selection and recombination rate play in determining the levels of genetic diversity and the efficiency of natural selection across the non-human homininae genomes remains unresolved. In humans, it was observed many years ago that levels of diversity at putatively neutral sites are correlated to the rate of recombination (Lercher & Hurst 2002; Hellmann et al. 2005). Since the rate of substitution is also correlated to the rate of recombination it seemed likely that at least part of the correlation between diversity and the rate of recombination was due to a mutagenic effect of recombination. There is now good evidence that recombination is mutagenic in humans (Pratto et al. 2014; Francioli et al. 2015; Arbeithuber et al. 2015; Halldorsson et al. 2019) and recent analyses of the correlation between diversity and the rate of mutation, as inferred from rates of *de novo* mutations in human trios, suggests that much, but not all, of the variation in diversity across the human genome can be explained by variation in the rate of mutation at the 100KB and 1MB scale (Smith et al. 2018). However, several lines of evidence suggest that selection at linked sites may also affect neutral and selected diversity across the human genome. First, it has been observed that levels of diversity are negatively correlated to gene density (Payseur & Nachman 2002). Second, levels of non-coding diversity are lower near functional DNA elements in humans and non-human primates (McVicker et al. 2009; Enard et al. 2014; Nam et al. 2017). Third, the rate of non-synonymous to synonymous substitution is positively correlated to gene (and exon) density (Bullaughey et al. 2008). Fourth, (Hussin et al. 2015) showed that exons in regions of low recombination are significantly enriched for deleterious and disease-associated variants consistent with variation in the intensity of selection at linked sites generating variation in the efficiency of purifying selection along the genome.

In our study, we find that the three genomic variables: recombination rate, mutation rate and the density of conserved sites are correlated to each other. We show that the levels of both putatively neutral non-coding and putatively selected non-synonymous variation are correlated to those genomic variables in most homininae, but that the relative importance of each genomic variable is different for non-coding and non-synonymous polymorphisms. Interestingly, we also find evidence that indicates variation in the efficiency of negative selection likely generated by interference among deleterious mutations in the genome of all the homininae. Finally, we find little impact of gBGC in our analyses and conclusions when interrogating only GC-conservative mutations.

## Materials and Methods

### Population genomic data

SNP calls from the autosomes were retrieved from (Prado-Martinez et al. 2013) for five great ape populations: *Homo sapiens, Pan paniscus, Pan troglodytes ellioti, Pan troglodytes verus* and *Gorilla gorilla gorilla.* Hereafter we refer to these species as humans, bonobos, Nigeria-Cameroon chimpanzees, western chimpanzees and gorillas, respectively. We analyzed 8 chromosomes per position in all populations (see details below). In Prado-Martinez et al. (2013) all reads were mapped to the human reference genome (hg18), but we used lift-over to hg19/GRCh37.75 coordinates to take advantage of more recent functional annotations (see below). To avoid errors introduced by mismapping due to paralogous variants and repetitive sequences, we also restrict all analyses to a set of sites with a unique mapping to the human genome (Cagan et al. 2016). Additionally, we require positions to have at least 5-fold coverage in all individuals per species. Only the resulting set of sites are used in further analyses (Supplementary Table 1). The final list of analyzed positions is available upon request.

### Genome annotation and identification of putatively neutral non-coding sites

Genomes were annotated using the SnpEff and SnpSift software (Cingolani et al. 2012) (version 4.3m, last accessed June 2017) and the human database GRCh37.75. We extracted 0-fold degenerate sites from the codon information provided by SnpEff (4-fold, 2-fold and 3-fold degenerate sites are discarded). We assume that the degeneracy and gene annotations are identical across species. In order to obtain putatively neutral non-coding sites, we applied stringent filters: (1) We kept sites annotated only as intronic or intergenic (splicing sites, UTRs, coding or transcribed non-coding genes are discarded). (2) We removed GERP elements (Davydov et al. 2010) and positions with a PhastCons score > 5% in primates and/or in 100 vertebrate species (Siepel et al. 2005). In this way, we removed conserved sites at different phylogenetic depths. Note that GERP elements were calculated in a multiple species alignment of the human genome to 33 other mammalian species (the most distant mammalian species is Platypus), while the PhastCons scores used here are based on a multiple species alignment of the human genome to 9 other primates (the most distant is bushbaby) and a multiple species alignment of the human genome to 99 other vertebrate species (the most distant is zebrafish). (3) Some sites might have become functional very recently. Thus, we removed *DNase* I hypersensitivity sites across multiple human tissues (Song et al. 2011) (downloaded from: http://ftp.ebi.ac.uk/pub/databases/ensembl/encode/integration_data_jan2011/byDataType/openchrom/jan2011/combined_peaks/). (4) Predicted transcription factor binding sites detected in humans were also excluded (Cingolani et al. 2012). (5) Hypermutable *CpG* sites in humans and the rest of species were excluded to remove variation in mutation rate due to variation in GC-content. This last filter was also applied to coding sites and mutation rate estimates (see below).

### Genomic windows and statistics

We split the autosomes in non-overlapping windows of 50KB, and for each window we estimate: (1) diversity at putatively neutral non-coding (NC) sites, (2) 0-fold degenerate (N) sites, (3) GC-conservative substitution rate at NC sites (our main proxy of the mutation rate), (4) the density of conserved sites, (5) the rate of recombination (RR, mainly crossing overs) and (6) the rate of *de novo* mutations (DNMs) (an alternative proxy of the mutation rate).

### Recombination maps and the density of conserved sites

Population recombination rate estimates for non-human great apes are retrieved from (Stevison et al. 2016) and (Auton et al. 2012) and human population recombination rates from HapMap (Myers et al. 2005). These recombination maps are LD-based representing both crossing over and gene conversion events. However, as gene conversion tracts tend to be very small in humans (50-150 bp) (Jeffreys & May 2004; Padhukasahasram & Rannala 2013), these maps measure mainly crossing over events. Blocks for each non-human genome that are syntenic with human were identified as in Stevison et al. (2016). To estimate the density of conserved sites we used as before GERP elements (Davydov et al. 2010) and PhastCons scores (Siepel et al. 2005), but this time we labeled and count all GERP elements and/or positions with a PhastCons score > 50% in primates and/or positions with a PhastCons score > 50% in 100 vertebrate species in a given 50KB window. We discarded the number of unsequenced nucleotides in the human reference genome to estimate the density of conserved sites.

### Polymorphism and mutation rate estimates

For a fair comparison between species, we downsampled our population genomic data to 8 haploid chromosomes per position. Positions called in less than 8 chromosomes were excluded. For each window we counted the total number of analyzable polymorphic sites (L_P,N_ and L_P,NC_) and the number of segregating sites (S_N_ and S_NC_) to get the Watterson estimator (θ, (Watterson 1975)) for NC and N sites, respectively. We did this for all point mutations and for GC-conservative point mutations.

For divergence estimates, we counted the total number of analyzable divergent non-coding sites (L_D,NC_) from a multiple species alignment between one randomly sampled Nigeria-Cameroon chimpanzee, western chimpanzee, bonobo, gorilla, and human chromosome. This multiple species alignment is generated from Prado-Martinez et al. (2013) original VCF file. Then, to estimate our proxy of the mutation rate (d_NC_) in each window, and given that there are few GC-conservative substitutions per window, we sum all GC-conservative substitutions (D_NC_) occurring in the homininae tree and divide it by L_D,NC_. Thus, our proxy of the mutation rate is the same for all species and it is unaffected by gBGC. Given that species specific substitution rates at 50KB are strongly correlated between species pairs (Supplementary Table 2) we believe this is a reasonable approach to gain statistical power without losing resolution. For some validation analyses, we also used the rate of DNMs per window in humans taking our DNMs from the studies of Jónsson et al. (2017), Wong et al. (2016) and Francioli et al. (2015) as an alternative proxy of the regional mutation rate. We only considered non-CpG DNMs but this time we considered both GC-conservative and non-GC-conservative DNMs.

### Hypergeometric sampling and grouping

We are interested in the effect of the mutation rate, the rate of recombination and the density of conserved sites on the efficiency of negative selection across the homininae genomes. We consider first the log(θ_N_/θ_NC_) and its relationship to the mutation rate, the rate of recombination and the density of conserved sites, and then its relationship to a measure of the local effective population size, log(θ_NC_/d_NC_). log(θ_N_/θ_NC_) and log(θ_NC_/d_NC_) are undefined if θ_N_, θ_NC_ or d_NC_ are zero, we, therefore, combined data across windows in the following manner. We split the number of non-coding polymorphisms (S_NC_) in each window into three parts using a hypergeometric distribution (see Supplementary Figure 1 for details). We used θ_NC,1_ to rank and bin windows into 50 groups. We then used θ_NC,2_ as our unbiased measure of the non-coding diversity and used θ_NC,3_ to estimate the ratio θ_N_/θ_NC_. We applied two methods to combine data across windows. In both methods we split S_NC_ into three statistically independent estimates using a hypergeometric distribution. In the first method, we include all windows that have non-coding sites, irrespective of whether they have coding sites. In the second method, we only include windows with coding sites. The second method yields about 43% of the data-points of the first method due to the requirement that windows have both coding and non-coding sites. The rationale for using two methods is that, for the non-coding analyses, conserved non-coding sites might be a source of selection at linked sites. We present results from method 1 for non-coding results and from method 2 for non-synonymous results. There are some outlier regions with very high θ_NC_ and d_NC_ values. These have a disproportionate influence over the statistics. We hypothesize that those high diversity regions might overlap with genes under balancing selection and/or low complexity repetitive regions. We excluded all windows overlapping with the MHC locus and other top candidate genes under balancing selection in great apes retrieved from (Azevedo et al. 2015). To remove low complexity regions we analyzed positions outside the DAC blacklisted regions (from: https://genome.ucsc.edu/cgi-bin/hgFileUi?db=hg19&g=wgEncodeMapability). This combined filtering resulted in the removal of all outlier regions and the exclusion of approximately 3% of the windows and 10% of the sites.

### Expected relationship between θ_N_/θ_NC_ and θ_NC_

θ_N_ and θ_NC_ are expected to be correlated through variation in the mutation rate and/or the *N_e_*. If we assume that the distribution of fitness effects (DFE) of new deleterious mutations follows a gamma distribution, then, under free recombination, the slope (*b*) of the relationship between θ_N_/θ_NC_ and θ_NC_ in a log-log scale informs us about the source of this variation (Welch et al. 2008). If there is no variation in *N_e_* and all variation in θ_N_ and θ_NC_ is due to variation in mutation rate, then we expect *b* = -1. In contrast, if all the variation in θ_N_ and θ_NC_ comes from variation in the *N_e_*, then we expect *b* = -*β* (Welch et al. 2008), where *β* is the shape parameter of the gamma distribution. Finally, if θ_N_ and θ_NC_ are independent we expect *b* = 0. Forward simulations with background selection (that is, limited recombination plus deleterious and neutral mutations) and variation in *N_e_* among loci have shown that the shape of the deleterious DFE can be successfully estimated with the slope between log(θ_N_/θ_NC_) and log(θ_NC_) after correcting for variation in the mutation rate (James et al. 2017). Castellano et al. (2018) showed that the slope is overestimated when HHs are not accounted for due to the faster recovery of the levels of deleterious variation compared to the levels of neutral variation. Recently, this relationship has been studied in more than 50 other species confirming that beneficial mutations must be invoked to explain the slope in large *N_e_* species (Chen et al. 2019).

### Statistical analyses

All statistical analyses were performed within the R framework (version 3.4.4). Here we want to explain how our dependent variable, the number of SNPs in a given window, which is discrete and over-dispersed data (Supplementary Figure 2), is related to our three genomic variables. To do that, we implemented a negative binomial regression by means of the R function *glm.nb* from the R package *MASS* (Venables & Ripley 2002). We modeled the log of the expected number of non-coding or non-synonymous SNPs as a function of the predictor variables; recombination rate, d_NC_, the density of conserved sites and the number of non-coding or 0-fold degenerate sites in a given 50KB window, respectively. We can interpret the negative binomial regression coefficient as follows: for one unit of change in the predictor variable, the difference in the logs of expected counts of the response variable is expected to change by the respective regression coefficient, given the other predictor variables in the model are held constant. To assess the relative importance of each genomic variable, we reported standardized regression coefficients to make variances of dependent and independent variables equal 1. These standardized coefficients refer to how many standard deviations the log of the expected number of SNPs will change, per standard deviation increase in the predictor variable. To explain the variation in our statistic of the efficiency of negative selection, log(θ_N_/θ_NC_), we used a standard multiple linear regression using the R function *lm*. As before, the standardized regression coefficients are used to assess the relative importance of each genomic variable. Finally, to estimate the variance inflation factor we used the R function *vif* from the package *car* (Fox & Weisberg 2018) and to perform the bivariate correlations we used the R function *cor.test* and the non-parametric Spearman method (Holland & Wolfe 1973).

## Results

### Genomic determinants of non-coding and non-synonymous diversity

We are interested in how genetic variation is distributed along the genomes of the homininae and in particular, the role that selection at linked sites and mutation rate variation might play in this distribution. To see whether these patterns are consistent across all homininae, we used the data of Prado-Martinez et al. (2013) sampling 8 random chromosomes per position in all five species and populations: humans, Nigeria-Cameroon chimpanzees, western chimpanzees, bonobos and gorillas. To allow an unbiased comparison with the other homininae we use the cosmopolitan human sample from the Great Apes Project. Supplementary Table 1 shows the summary statistics of our analyzed dataset. We use the same density of conserved sites and mutation rate for all species (see Supplementary Table 2 for the correlation of lineage specific substitution rates between pairs of species), while the estimates of recombination rate are population-specific and come from publicly available recombination rate maps (Stevison et al. 2016; Auton et al. 2012; Myers et al. 2005). We estimate the level of diversity at non-coding sites and 0-fold degenerate sites in 50KB windows across the autosomes. We exclude non-coding sites that are inferred to be subject to natural selection (based on the conservation of sites across species and other potentially functional annotations such coding and non-coding genes, UTRs, *DNase I* hypersensitivity sites and transcription factor binding sites).

We expect the level of genetic diversity at both selected and neutral sites to depend on the mutation rate, the rate of recombination and the density of conserved sites, because each of these factors are expected to affect the diversity either directly, or indirectly. To estimate the mutation rate there are two options: using *de novo* mutations (DNMs) that have been discovered by the sequencing of trios or using the divergence between species. Neither of these methods is perfect. We currently have too few DNMs to estimate the mutation rate reliably at the 50KB scale and attempts to predict the mutation rate of DNMs based on genomic features have so far proved to be unreliable (Smith et al. 2018). The divergence between species is also not a completely satisfactory measure of the mutation rate either, for several reasons. We have used GC-conservative substitutions (i.e. A<>T and G<>C), since these are not affected by gBGC, a process known to affect substitution rates (Duret & Arndt 2008; Smith et al. 2018), and the rate of different types of mutation appear to be strongly correlated at the 100KB and 1MB scales, suggesting that GC-conservative mutations should therefore be a reasonable measure of the overall mutation rate (Smith et al. 2018). However, the mutation rate appears to evolve at large scales (Terekhanova et al. 2017; Smith et al. 2018), and some of the variation in the substitution rate is due to variation in the depth of the genealogy in the ancestors of the homininae (Phung et al. 2016).

The rate of recombination, the density of selected sites and our measure of the mutation rate are correlated to each other (Table 1). We confirm the positive correlation between the mutation rate and the rate of recombination in humans but also, for the first time, in the other non-human homininae (Table 1; Supplementary Table 3). We also find a negative correlation between the density of conserved sites and the rate of recombination in humans (as in (McVean et al. 2004; Kong et al. 2010)) and the rest of homininae (Table 1; Supplementary Table 3). We find that this negative correlation is driven by conserved coding sites but not by conserved non-coding sites in agreement with the lower rate of recombination seen in exons and nearby non-coding regions (Supplementary Analyses). Interestingly, there is a strong negative correlation between the density of conserved sites and our measure of the mutation rate. We investigated this further in humans using three large publicly available *de novo* mutation (DNM) datasets from trios (Francioli et al. 2015; Wong et al. 2016; Jónsson et al. 2017) (Supplementary Analyses). We find that the density of putatively neutral DNMs is either significantly positively correlated or uncorrelated to the density of conserved sites, depending on which dataset of DNMs is considered. Hence, we do not know if the negative correlation between our measure of the mutation rate, d_NC_, and the density of conserved sites is a consequence of co-variation in the mutation rate and the density of conserved sites, or whether d_NC_ is reduced in regions with high density of conserved sites (despite our stringent filtering, we might not be masking all sites under purifying selection).

**Table 1.**
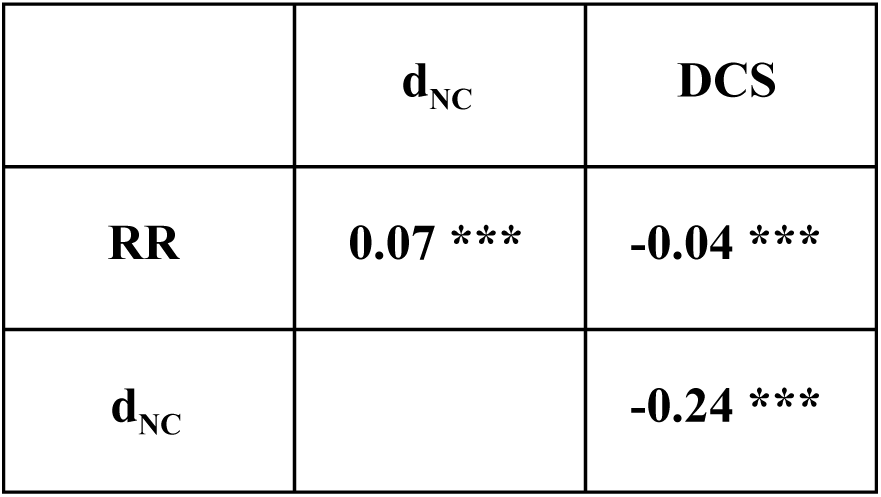
Spearman correlations between genomic variables at 50 KB in humans. RR: Recombination rate. DCS: density of conserved sequences. d_NC_: GC-conservative substitution rate at non-coding sites (mutation rate proxy).

As expected, we find that non-coding diversity is positively correlated to the rate of recombination and our measure of the mutation rate, and negatively correlated to the density of selected sites if we run a bivariate analysis (Supplementary Figure 3-7). In a negative binomial multiple regression all three genomic variables are significant (Figure 1A; Supplementary Table 4A) (note that variance inflation factors are close to 1 for each of our genomic variables suggesting that collinearity is not a problem). Standardized regression coefficients suggest that all three genomic variables are equally important in determining levels of putatively neutral diversity in humans and non-human homininae (Figure 1A; Supplementary Table 4A). The only exception is western chimpanzees which show a weak correlation between non-coding diversity and the rate of recombination, but a stronger correlation between non-coding diversity and mutation rate.

**Fig. 1.**
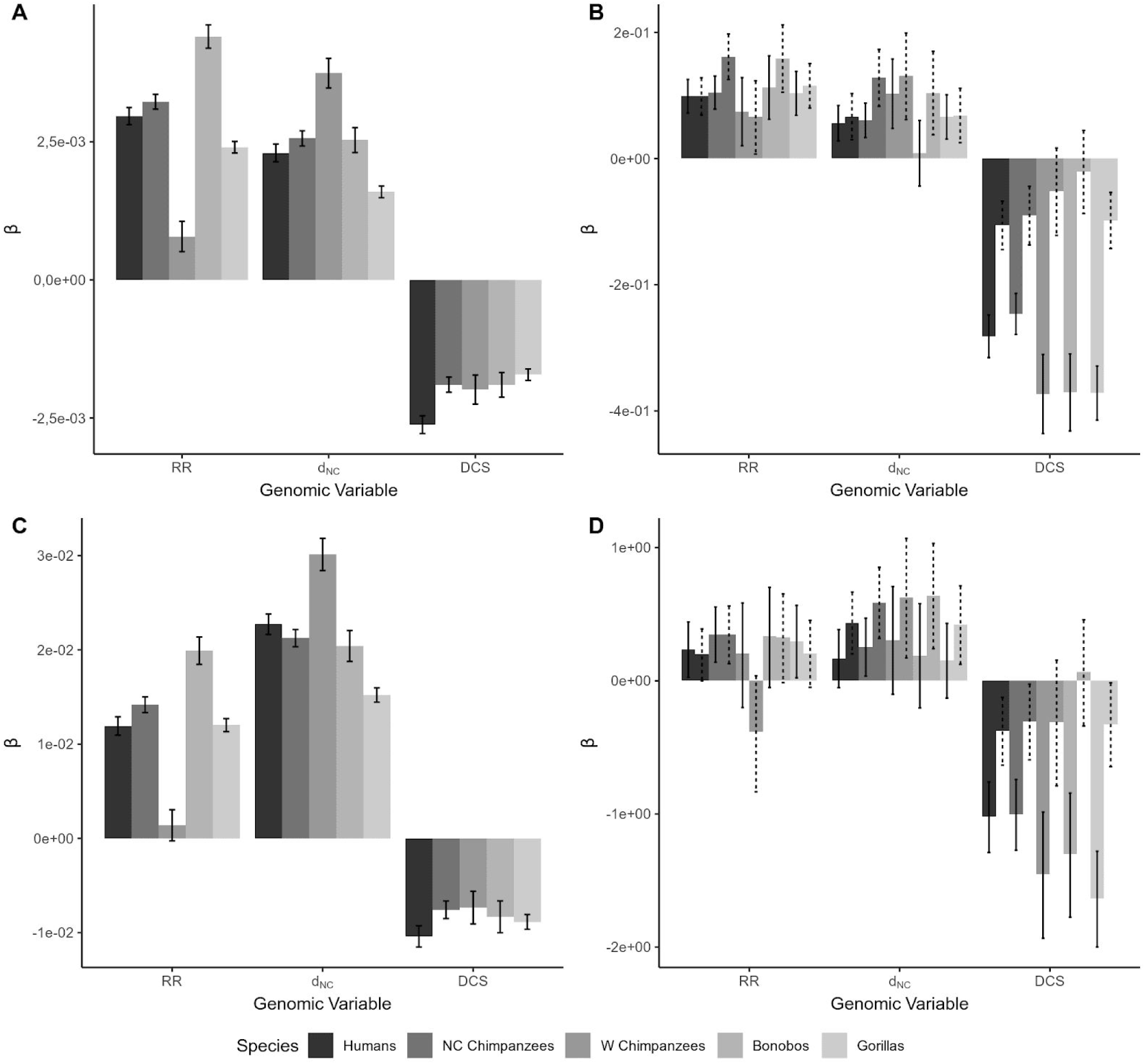
The standardized regression coefficients (*β*) in the y-axis and the three genomic variables (RR: recombination rate, d_NC_: mutation rate and DCS: density of conserved sequences) in the x-axis for each species. All non-coding mutations (A), all non-synonymous mutations with the respective matching set of (downsampled) non-coding mutations (B), GC-conservative non-coding mutations (C), GC-conservative non-synonymous mutations with the respective matching set of (downsampled) non-coding mutations (D). The solid error bars indicate the confidence intervals 95% of the original dataset and the dashed error bars represent the confidence intervals 95% of the downsampled non-coding dataset for comparison.

The results for 0-fold degenerate sites are qualitatively consistent with the non-coding results; non-synonymous diversity is positively correlated to the rate of recombination and mutation, and negatively correlated to the density of selected sites. Here, all factors remain significant in most species in the negative binomial multiple regression (Figure 1B; Supplementary Table 4B). The density of conserved sites is the strongest correlate as judged by standardized regression coefficients; it has more than twice the impact on non-synonymous diversity as the rate of recombination. Our measure of the mutation rate has slightly more than half of the impact of recombination rate. These patterns are consistent across most of the homininae. However, the results for 0-fold degenerate sites are quantitatively different from that observed for non-coding sites, in which diversity is equally strongly correlated to the three genomic factors. This difference between non-synonymous and non-coding sites does not appear to be due to differences between the windows that have non-synonymous sites and those that do not. If we only use windows that have non-synonymous sites and down-sample the number of non-coding polymorphisms so that it matches the number of non-synonymous SNPs on average we find that non-coding diversity is equally strongly correlated to each of the genomic variables and the density of conserved sites is not the dominant correlate (Figure 1B). Our results therefore suggest that there are genuine differences in the relative impact of these three genomic variables on non-synonymous and non-coding diversity. Finally, given the collinearity between explanatory variables (Table 1) we computed the variance inflation factor (VIF) of the models used to generate Figure 1 (and Supplementary Table 4). VIF values are all close to one suggesting that multicollinearity is not an issue with this analysis.

### Biased gene conversion

The correlation between the rate of recombination and diversity might be (directly) mediated by GC-biased gene conversion (gBGC). We repeated our analyses restricting the analysis to GC-conservative SNPs. As expected, our results are largely unaffected by this restriction; non-coding diversity remains correlated to all factors (Figure 1C; Supplementary Table 5A). For non-synonymous diversity, we again find qualitatively similar results to those using all mutations, although the correlation between non-synonymous diversity and our estimate of the mutation is generally non-significant (Figure 1D; Supplementary Table 5B).

To investigate in more detail whether gBGC has an effect on the correlations between diversity and our three genomic variables, we considered the correlation between diversity and recombination rate after down-sampling the non-GC-conservative mutations to yield the same average number of mutations per 50KB as we find in the GC-conservative dataset. GC-conservative mutations represent 15-18% of all SNPs in the homininae subfamily (Supplementary Table 1). Although the effect is small (Figure 2A), the correlation between θ_NC_ and the rate of recombination is greater for non-GC-conservative mutations than GC-conservative mutations in all homininae, despite the fact that there are fewer GC-conservative mutations; the consistency across the homininae is significant as judged by a sign test (*P*-value < 0.01). This suggests that gBGC has a significant but small effect on the correlation between recombination rate and non-coding diversity. The strength of the correlation between θ_N_ and the rate of recombination is again similar between mutation types and the difference is only significant in humans (*P*-value = 0.043) (Figure 2B).

**Fig. 2.**
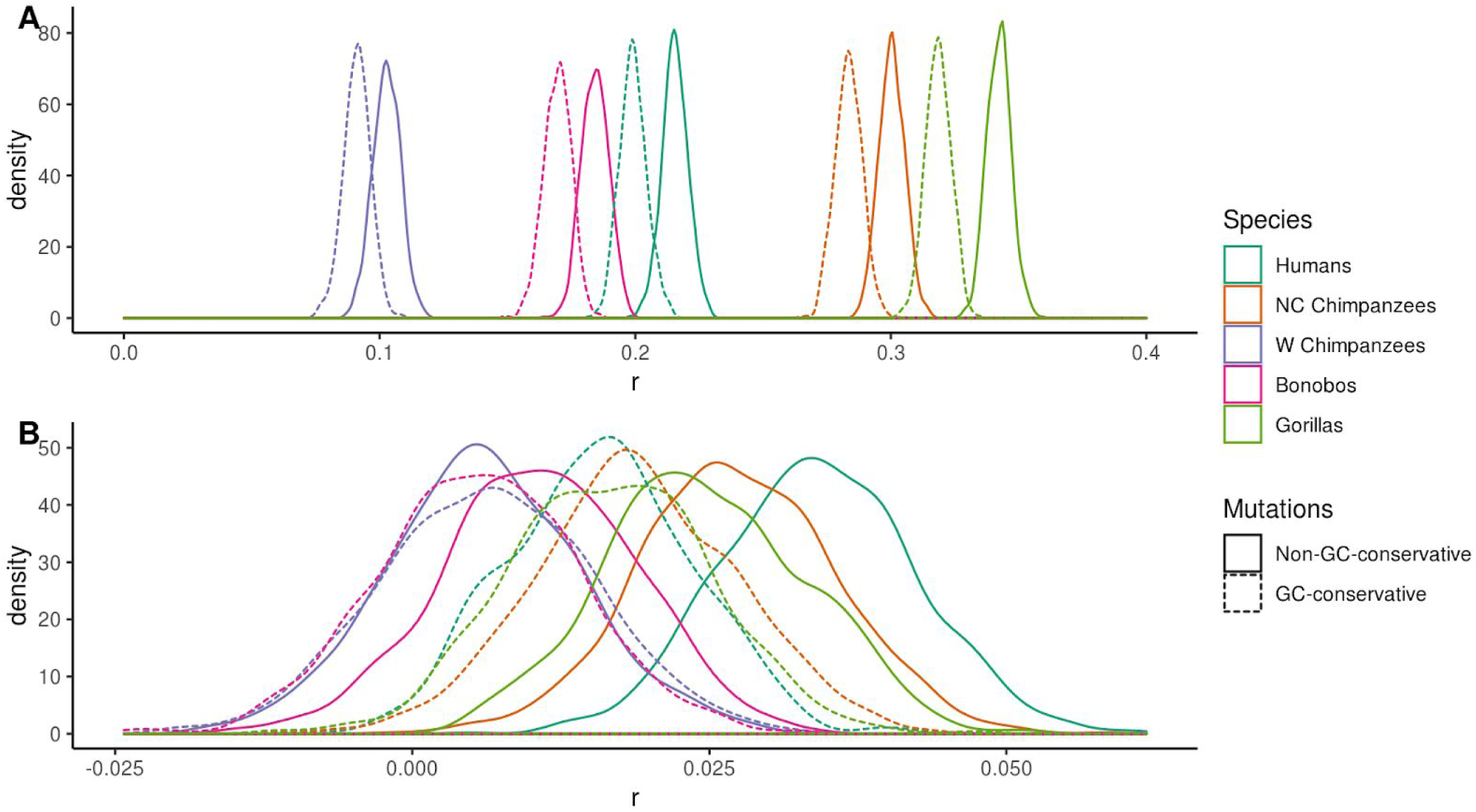
Relative effect of gBGC on the relationship between recombination rate and non-coding diversity (A) and non-synonymous diversity (B). Distribution of the Spearman rank correlation coefficients (*r*) across 1000 bootstrap replicates for non-GC-conservative mutations (downsampled to match GC-conservative diversity) and GC-conservative mutations.

### Efficiency of negative selection

Finally, we sought to investigate whether there is an influence of selection at linked sites on the efficiency of negative selection along the genomes by considering whether θ_N_/θ_NC_ is correlated to the recombination rate, d_NC_ and the density of conserved sites. If there is variation in the effects of selection at linked sites across the homininae genome then we would expect θ_N_/θ_NC_ to be negatively correlated to recombination rate, and positively correlated to the mutation rate and the density of conserved sites. Because we have very few coding sites in each window, and hence very few non-synonymous SNPs, we grouped windows together into 50 groups that each contains on average ∼1800 non-coding SNPs, and ∼25 non-synonymous SNPs (in humans). Using a hypergeometric sampling (Supplementary Figure 1), we grouped windows by splitting the non-coding SNPs in each window into three independent estimates of the non-coding diversity; we used one estimate to rank and group windows, the other as our unbiased estimate of the diversity in a group of windows and the third to estimate the ratio θ_N_/θ_NC_. We find that θ_N_/θ_NC_ is generally negatively correlated to the rate of recombination and our estimate of the mutation rate, d_NC_, and positively correlated to the density of conserved sites, when the correlations are performed individually (Table 2).

**Table 2.**
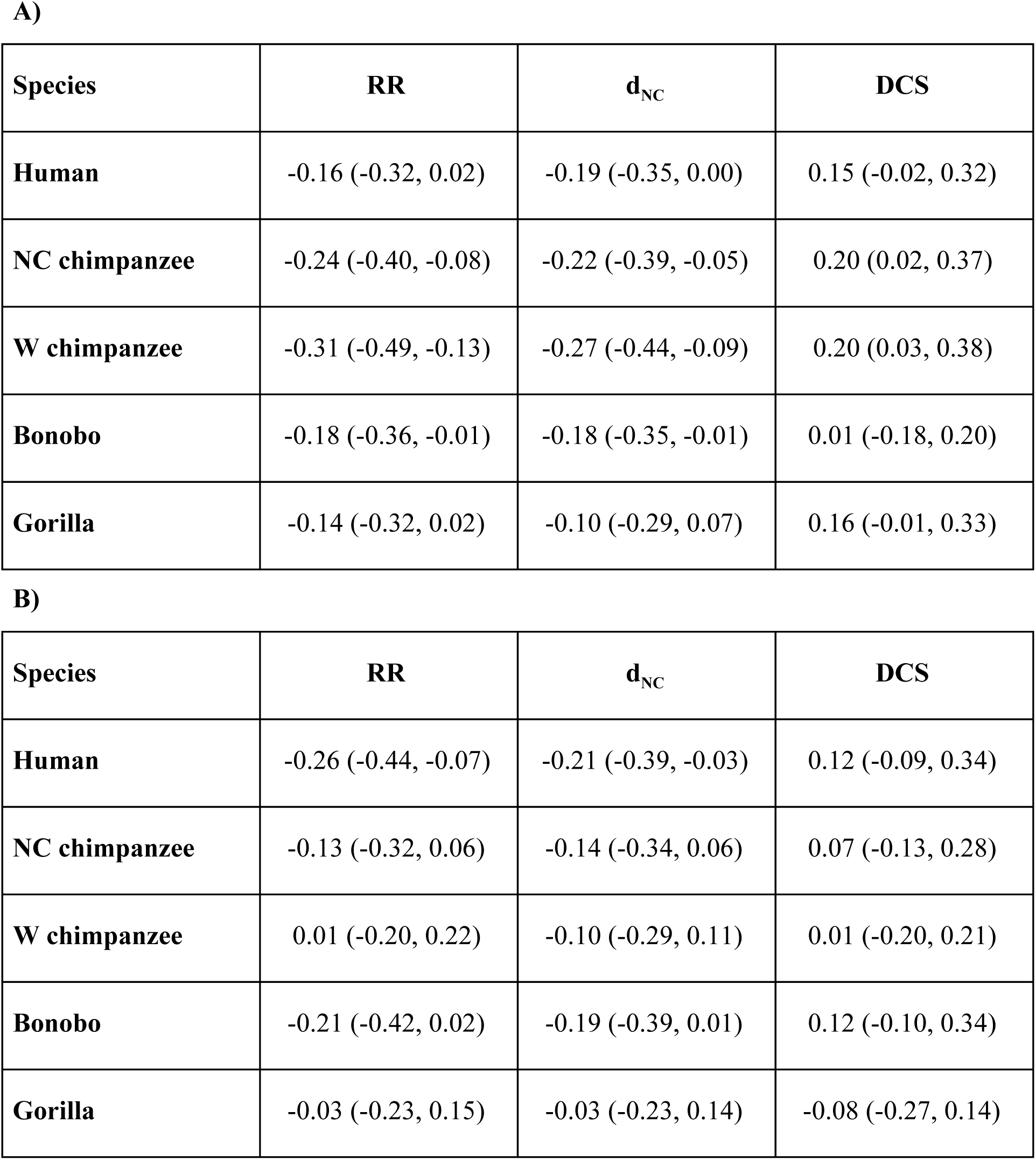
The Spearman rank correlation coefficients of θ_N_/θ_NC_ versus RR, d_NC_ and the density of conserved sites (DCS) A) for all SNPs and B) for GC-conservative SNPs. Given in parentheses are the 95% confidence intervals by bootstrapping.

| Species | RR | $d_{NC}$ | DCS |
| --- | --- | --- | --- |
| **Human** | -0.16 (-0.32, 0.02) | -0.19 (-0.35, 0.00) | 0.15 (-0.02, 0.32) |
| **NC chimpanzee** | -0.24 (-0.40, -0.08) | -0.22 (-0.39, -0.05) | 0.20 (0.02, 0.37) |
| **W chimpanzee** | -0.31 (-0.49, -0.13) | -0.27 (-0.44, -0.09) | 0.20 (0.03, 0.38) |
| **Bonobo** | -0.18 (-0.36, -0.01) | -0.18 (-0.35, -0.01) | 0.01 (-0.18, 0.20) |
| **Gorilla** | -0.14 (-0.32, 0.02) | -0.10 (-0.29, 0.07) | 0.16 (-0.01, 0.33) |

**Table 2.** The Spearman rank correlation coefficients of $\theta_N/\theta_{NC}$ versus RR, $d_{NC}$ and the density of conserved sites (DCS) A) for all SNPs and B) for GC-conservative SNPs. Given in parentheses are the 95% confidence intervals by bootstrapping.
| Species | RR | $d_{NC}$ | DCS |
| --- | --- | --- | --- |
| **Human** | -0.26 (-0.44, -0.07) | -0.21 (-0.39, -0.03) | 0.12 (-0.09, 0.34) |
| **NC chimpanzee** | -0.13 (-0.32, 0.06) | -0.14 (-0.34, 0.06) | 0.07 (-0.13, 0.28) |
| **W chimpanzee** | 0.01 (-0.20, 0.22) | -0.10 (-0.29, 0.11) | 0.01 (-0.20, 0.21) |
| **Bonobo** | -0.21 (-0.42, 0.02) | -0.19 (-0.39, 0.01) | 0.12 (-0.10, 0.34) |
| **Gorilla** | -0.03 (-0.23, 0.15) | -0.03 (-0.23, 0.14) | -0.08 (-0.27, 0.14) |

The unexpected negative correlation with d_NC_ might be due to the positive correlation between the rate of recombination and d_NC_ (Table 1; Supplementary Table 3). In fact, none of these factors remain significant when we perform a multiple regression (Supplementary Table 6). The reason may be strong correlations between recombination rate, d_NC_ and density of conserved sites when we group windows introducing multicollinearity. Indeed, the variance inflation factor (VIF) is > 4.5 for each variable.

Recombination rate, our measure of the mutation rate and the density of conserved sites are likely to be crude predictors of the effects of selection at linked sites compared to the realized level of neutral diversity in a given window, after accounting for mutation rate variation. We, therefore, investigated whether θ_N_/θ_NC_ is correlated to a measure of the effective population size (*N_e_*) of a window, estimated by dividing the non-coding diversity by our estimate of the mutation rate: i.e. θ_NC_/d_NC_. We find that θ_N_/θ_NC_ is significantly negatively correlated to our measure of the local *N_e_* in all homininae species (Figure 3). The slope of this relationship on a log-log scale is expected to equal to the negative value of the shape parameter of the distribution of fitness effects, if the distribution is gamma distributed (Welch et al. 2008). The slope of this relationship in humans is -0.2 which is consistent with estimates of the distribution made from the site frequency spectrum (Eyre-Walker et al. 2006; Boyko et al. 2008; Eyre-Walker & Keightley 2009; Kim et al. 2017). The other non-human homininae show similar slopes (Table 3) suggesting that, in these species, the shape of the distribution of fitness effects for new deleterious mutations is quite stable (as recently reported by (Castellano et al. 2019) using the site frequency spectrum). Due to the lack of statistical power when we consider GC-conservative mutations the slope estimates become very noisy. The slope is not significantly different from 0 in the two chimpanzee populations and gorillas and not significantly different from 1 in humans and bonobos.

**Fig. 3.**
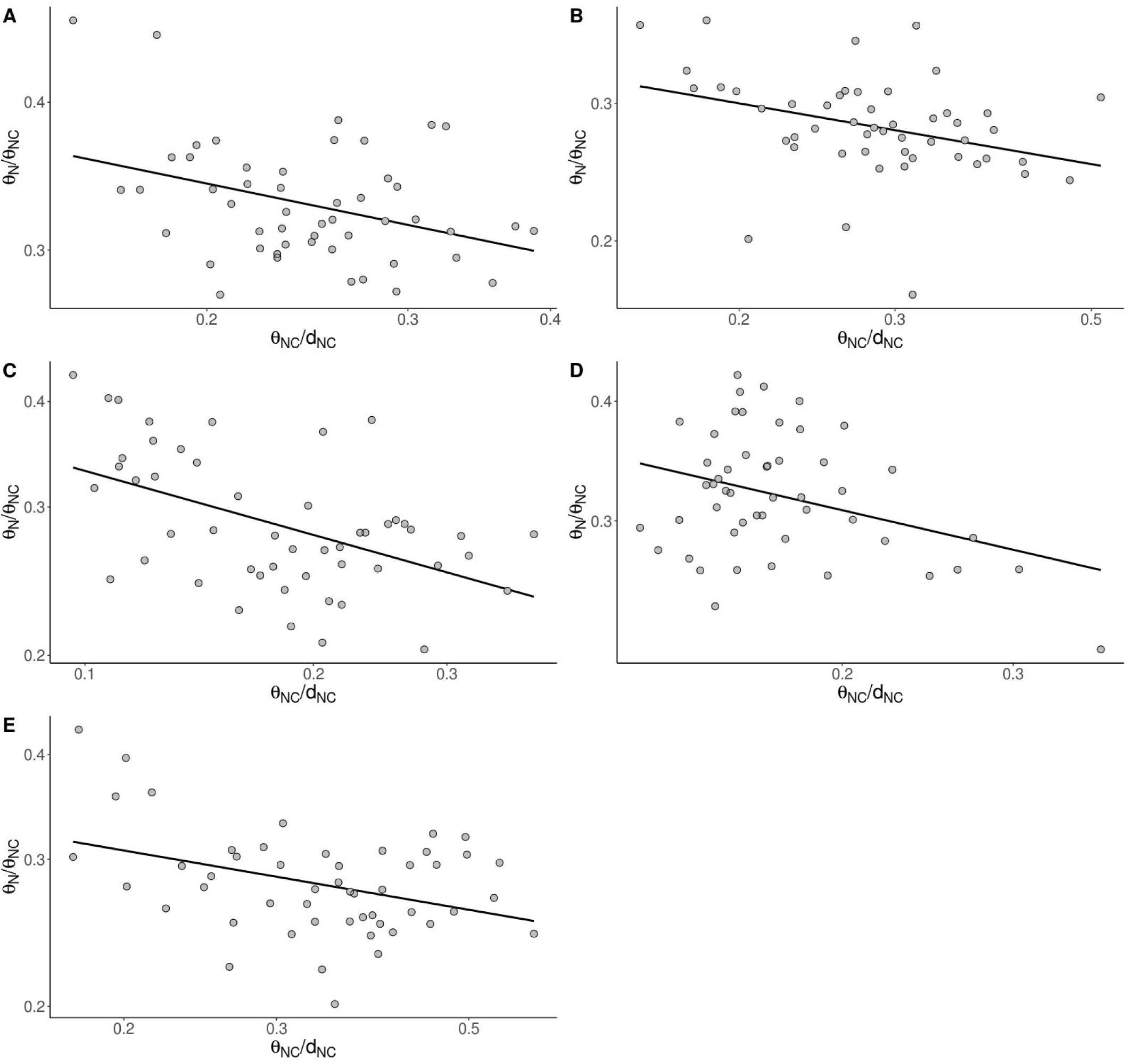
Relationship between θ_N_/θ_NC_ and θ_NC_/d_NC_ in a log-log scale for all mutations in A) humans (*P*-value < 0.01), B) Nigeria-Cameroon chimpanzees (*P*-value < 0.05), C) western chimpanzees (*P*-value < 0.001), D) bonobos (*P*-value < 0.05) and E) gorillas (*P*-value < 0.01). Results grouping 50KB windows into 50 bins by non-coding diversity.

**Table 3.** Mean slope (b) of log(θ_N_/θ_NC_) ∼ log(θ_NC_/d_NC_) linear regression for all SNPs and GC-conservative SNPs. Given in parentheses are the 95% confidence intervals by bootstrapping. Results grouping 50KB windows into 50 bins by non-coding diversity.

| Species | All mutations | GC-conservative mutations |
| --- | --- | --- |
| **Human** | -0.20 (-0.31, -0.08) | -0.59 (-1.09, -0.13) |
| **NC chimpanzee** | -0.20 (-0.33, -0.07) | -0.17 (-0.60, 0.22) |
| **W chimpanzee** | -0.21 (-0.31, -0.11) | -0.04 (-0.43, 0.35) |
| **Bonobo** | -0.31 (-0.46, -0.15) | -0.71 (-1.20, -0.18) |
| **Gorilla** | -0.14 (-0.24, -0.03) | -0.05 (-0.38, 0.29) |

## Discussion

We have investigated the three genomic variables known to affect the level of genetic diversity across the genomes of humans and other homininae. We find highly significant effects of recombination rate, mutation rate and the density of conserved elements on levels of putatively neutral genetic diversity in most species, even if we restrict the analysis to GC-conservative polymorphisms, and hence remove the effects of gBGC. The positive correlation between diversity and our measure of the mutation rate is not surprising given that we expect regions of the genome with high rates of mutation to have high levels of diversity. Previous analyses have suggested that much of the variation in diversity at the 100KB level can be explained in terms of variation in the mutation rate (Smith et al. 2018). Non-coding diversity is also positively correlated to the rate of recombination and negatively correlated to the density of conserved sites. Similarly, the level of non-synonymous diversity for all mutations as well as for GC-conservative mutations is strongly negatively correlated to the density of conserved sites and positively correlated to the recombination rate and our measure of the mutation rate in most homininae. However, the pattern of correlation differs between the two sets of sites; whilst non-coding diversity is approximately equally strongly correlated to all three genomic variables, non-synonymous diversity is much more strongly correlated to the density of selected sites.

The strong negative correlation between non-synonymous diversity and the density of conserved sites may be explained if the coding mutation rate and the density of conserved sites co-vary along the genome. To explore this hypothesis, we compiled *de novo* mutation (DNM) data from three studies that have discovered large numbers of DNMs in the sequencing of trios (Supplementary Analyses). We then investigated the relationship between putatively selected DNMs and the density of conserved sites. We find a significantly negative correlation between putatively selected DNMs and the density of conserved sites in one DNM dataset (Jónsson et al. 2017), a significantly positive correlation in another dataset (Wong et al. 2016), and no correlation in the third dataset (Francioli et al. 2015). Such contradictory patterns have been observed before for other genomic variables and likely arise as a consequence of biases in the ascertainment of DNMs (Smith et al. 2018). Thus, it is not possible to conclude that the lower coding mutation rate in functionally rich regions is driving our strong negative correlation between θ_N_ and the density of conserved sites. There is a second, non-mutually exclusive interpretation to explain the importance of the density of conserved sites on the levels of non-synonymous diversity. That is co-variation of the DFE of new amino acid mutations and gene density along the genome. In other words, if genes in highly gene-dense regions are more constrained (their DFE has a lower proportion of new nearly neutral and slightly deleterious mutations and a greater proportion of new strongly deleterious mutations) than “isolated” genes, then this will explain the negative correlation between the gene density and θ_N_ without invoking selection at linked sites or a lower mutation rate in functionally rich regions. The degree to which gene density (plus the associated regulatory sequences) and the DFE co-vary is therefore an interesting question for further investigation. However, the significantly negative slope between log(θ_N_/θ_NC_) ∼ log(θ_NC_/d_NC_) (see below) suggests that selection at linked sites is indeed reducing the efficiency of purifying selection across the homininae genome. A third explanation may be the lower recombination rate reported at exons bodies (Kong et al. 2010; McVean et al. 2004; Myers et al. 2005). Hill-Robertson interference (Hill & Robertson 1966) will be then very intense at coding rich regions due to the concomitant reduction of recombination.

It should be appreciated that the correlations between recombination rate, the density of conserved sites and genetic diversity may be an artifact of deficiencies in multiple regression. Recombination is known to be mutagenic in humans (Lercher & Hurst 2002; Hellmann et al. 2005; Pratto et al. 2014; Francioli et al. 2015; Arbeithuber et al. 2015; Halldorsson et al. 2019) and since d_NC_ is an imperfect measure of the mutation rate, this may allow the rate of recombination to remain significant in a multiple regression. Similarly, there is a negative correlation between the density of conserved sites and our measure of the mutation rate, d_NC_. This could be due to regions of the genome with a high density of conserved sites having lower mutation rates, or due to the fact that we have not successfully masked all sites subject to selection (i.e. regions of the genome with high density of conserved sites might have a higher density of other sites subject to recent selection, and these sites will decrease d_NC_). To explore this possibility, we have also investigated the relationship between the density of putatively neutral DNMs in humans and the density of conserved elements. Again, we find that the density of DNMs at putatively neutral sites is either significantly positively or uncorrelated to the density of conserved sites, depending on which dataset of DNMs is considered (Supplementary Analyses). As a consequence, we do not know whether the negative correlation between d_NC_ and density of conserved sites is a consequence of variation in the mutation rate with the density of conserved sites, or that d_NC_ is reduced in regions with a high density of conserved sites because there are sites subject to selection that we have not masked.

However, we find additional evidence that selection at linked sites affects the efficiency of purifying selection. First, we find that our measure of the efficiency of purifying selection, θ_N_/θ_NC_, is negatively correlated to our estimate of the rate of recombination and positively correlated to the density of conserved sites in a bivariate analysis. Unfortunately, when we group data we find that our genomic variables are strongly correlated to each other and so it is not possible to disentangle which variable is actually correlated to θ_N_/θ_NC_ – i.e. there is a problem of multicollinearity in our multiple regression. Second, we find that θ_N_/θ_NC_ is negatively correlated to a measure of the effective population size of a window, θ_NC_/d_NC_. We find that the slope of the relationship in a log-log scale is consistent with the estimated shape parameter of the distribution of fitness effects in great apes (assuming the distribution is gamma distributed) (Castellano et al. 2019). This is in contrast to what has been observed in *Drosophila melanogaster*, in which the slope of the relationship between log(θ_N_/θ_S_) versus log(θ_S_) is significantly steeper than expected given an estimate of the DFE estimated from the site frequency spectrum; a similar pattern is apparent between species (Chen et al. 2017; James et al. 2017). Castellano et al. (2018) considered a number of explanations for this and concluded that it was most likely due to genetic hitch-hiking; they showed by simulation that hitch-hiking increases the slope of the relationship, because deleterious genetic variation recovers more rapidly after a hitch-hiking event than neutral genetic variation (Gordo & Dionisio 2005; Pennings et al. 2014; Brandvain & Wright 2016; Chen et al. 2019; Do et al. 2015). This is consistent with the high rates of adaptive evolution observed in *Drosophila* (Smith & Eyre-Walker 2002; Andolfatto 2005; Eyre-Walker & Keightley 2009; Enard et al. 2014). Rates of adaptive evolution seem to be lower in humans than in *Drosophila* (Gossmann et al. 2012; Galtier 2016; Rousselle et al. 2019) which is therefore consistent with the fact that the slope is similar to the shape parameter of the gamma distribution recently estimated across great apes (Castellano et al. 2019). However, note that (Zhen et al. 2018) have reported higher rates of adaptive substitutions in the past in humans than in *Drosophila* when accounting for ancestral population size. Hence, it is likely that most of the current variation in θ_N_/θ_NC_ along these genomes reflects variation in the efficiency of selection caused by selection at linked sites, mainly driven by weakly deleterious mutations.

Finally, we would like to highlight that interference among slightly deleterious mutations can potentially lead to indirect selection on recombination modifiers (Otto & Lenormand 2002; Bullaughey et al. 2008; Coop & Przeworski 2007) and that such selection may contribute to the evolution of fine-scale recombination patterns.

## Conclusions

We show that non-coding and non-synonymous diversity are positively correlated to both mutation rate and recombination rate, while the density of conserved sites is associated with low levels of genetic diversity in all homininae. Nonetheless, we also find genuine differences in the relative effect of these three genomic variables on the levels non-synonymous and non-coding diversity which deserve further investigation. The positive correlation with the rate of recombination and the negative correlation with the density of conserved sites is consistent with variation in the intensity of selection at linked sites along the genome. While the positive correlation with the mutation rate just indicates that the higher the number of new mutations per generation the higher the level of genetic diversity. We find a negative correlation between the ratio of the number of non-synonymous to non-coding polymorphisms and a measure of the effective population size across the homininae genome which suggests pervasive interference selection, mainly, among weakly deleterious variants.

## Supporting information

Supplementary Analyses

## Acknowledgments

We thank D. Weghorn and three anonymous reviewers for comments. We also thank Alex Cagan for sharing the set of sites with a unique mapping to the human genome with at least five-fold coverage in all individuals per species. This work was supported by the Danish Council For Independent Research (grant number 4181-00358).

