## Supplementary Analyses for "Impact of mutation rate and selection at linked sites on DNA variation across the genomes of humans and other homininae"

### Supplementary Material

#### Supplementary Analyses

##### *RR and the DCS*

Fine-scale (pedigree-based and LD-based) recombination maps have shown that RR is lower within exon bodies and the surrounding non-coding sites, around 10-30KB upstream and downstream exons (Kong et al. 2010; McVean et al. 2004; Myers et al. 2005). Interestingly, the first broad-scale (3MB) pedigree-based recombination map in humans found a positive correlation between RR and gene density (Kong et al. 2002). As suggested by McVean et al. (2004) this contradictory result can be explained if recombination tends to occur in regions of open chromatin (Nicolas 1998) and if recombination is mutagenic (Lercher & Hurst 2002; Hellmann et al. 2005; Smith et al. 2018). Hence, either new mutations (and recombination events) are generated within exon bodies but then they are purged by negative selection or mutation rate (and RR) is truly lower within and around exon bodies (below we further explore this possibility). We wonder whether the density of conserved non-coding sites or coding sites is more strongly correlated to RR. We find that the negative correlation is stronger with the density of conserved coding sites ($r$ = -0.1***) than with the density of conserved non-coding sites (those include potential regulatory elements, like transcription factor binding sites, UTRs, microRNAs and long non-coding genes along with metabolic RNAs such as tRNAs and rRNAs) which is non-significant ($r$ = -0.01) and the DCS (which can include both coding and non-coding sequences) (Table 1). This suggests that RR is only reduced (or apparently reduced) in coding regions and nearby regions but not in functional non-coding sequences. We obtain equivalent results whether we use the high-resolution pedigree map from Kong et al. (2010) or our LD-based measure of the RR (Myers et al. 2005) (data not shown).

##### *DNMs and the DCS*

We want to explore whether the rate of DNMs (Francioli et al. 2015; Wong et al. 2016; Jónsson et al. 2017) co-varies with the DCS as suggested by the negative correlation between our substitution-based measure of the mutation rate, $d_{NC}$, and the DCS (Table 1; Supplementary Figure 8G). A negative correlation between the rate of DNMs and the DCS might be driven by new detrimental mutations causing miscarriage (including here cases of stillbirth), by a greater DNA repair efficiency and/or by a lower generation of new mutations in functionally rich regions. To disentangle these effects, we split non-CpG DNMs and callable sites into two classes: putatively neutral and putatively selected (following the same filtering criteria used to get putatively neutral non-coding sites, see Materials and Methods). Note that while our definition of neutral change is very stringent, our definition of putatively selected mutation/site is not. All positions which are not cataloged as neutral are automatically labeled as selected (57%), this is obviously incorrect (Graur 2017). In Supplementary Figure 9 we show that the average rate of DNMs is ~1.7 times higher in our neutral category than in the selected category, this result is consistent across all datasets (Wilcoxon test $P$-value $< 0.001$). Thus, either extreme purifying selection is acting upon new mutations and/or the mutation rate is intrinsically higher in our set of putatively neutral sites. Supplementary Figure 8 shows the relationship between the rate of putatively neutral and putatively selected DNMs and the DCS for each DNM dataset. Hence, given the disparity of results across datasets we can not discard that our measure of the mutation rate, $d_{NC}$, decreases as the DCS increases due to the presence of unidentified sites under recent selection (probably lineage-specific) and that the negative correlation between genetic diversity and the DCS is driven by a lower mutation rate in functionally rich regions.
