## Supplementary Figures for "Impact of mutation rate and selection at linked sites on DNA variation across the genomes of humans and other homininae"

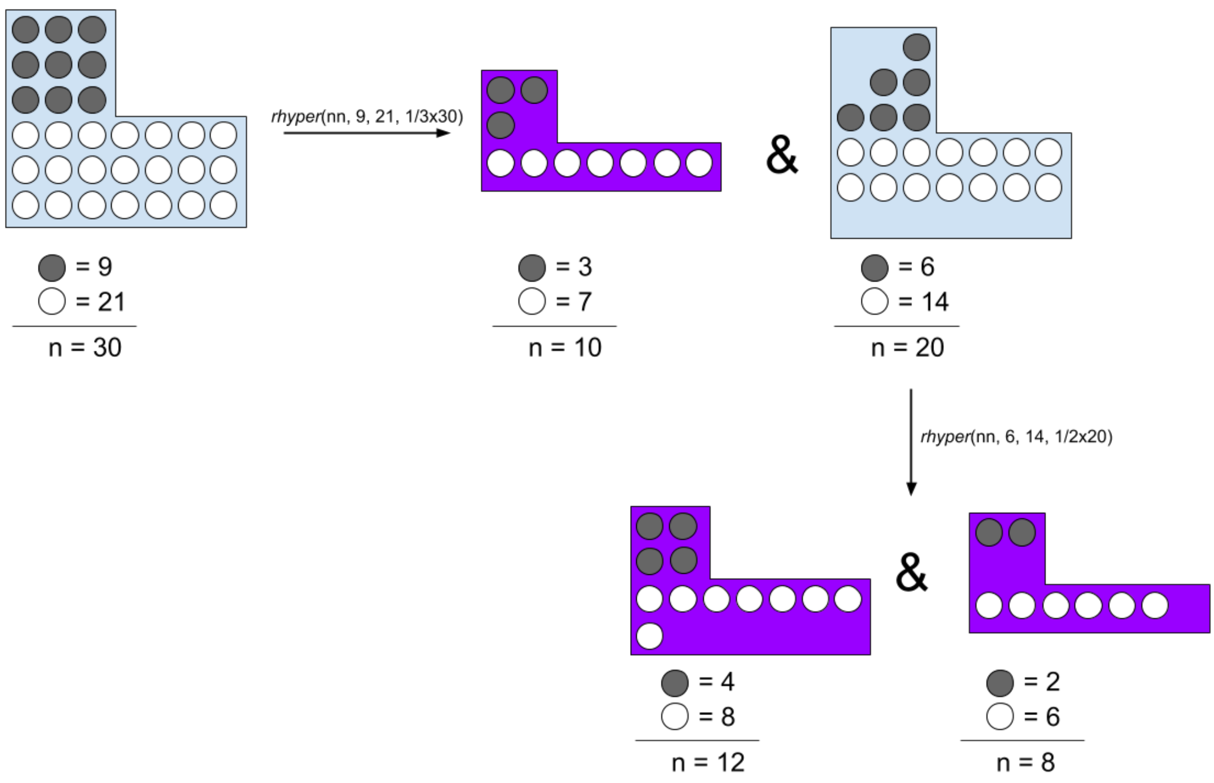


**Supplementary Figure 1**. Hypergeometric distribution approach to make three statistically independent samples (in purple) from discrete count data. Grey circles represent polymorphic sites and white circles represent monomorphic sites. *rhyper* is the R function used to generate the samples. *nn* is the number of draws, we did two draws per window, the first to obtain the first sample (n = 10), and the second to obtain the two other samples (n = 12 and n = 8).

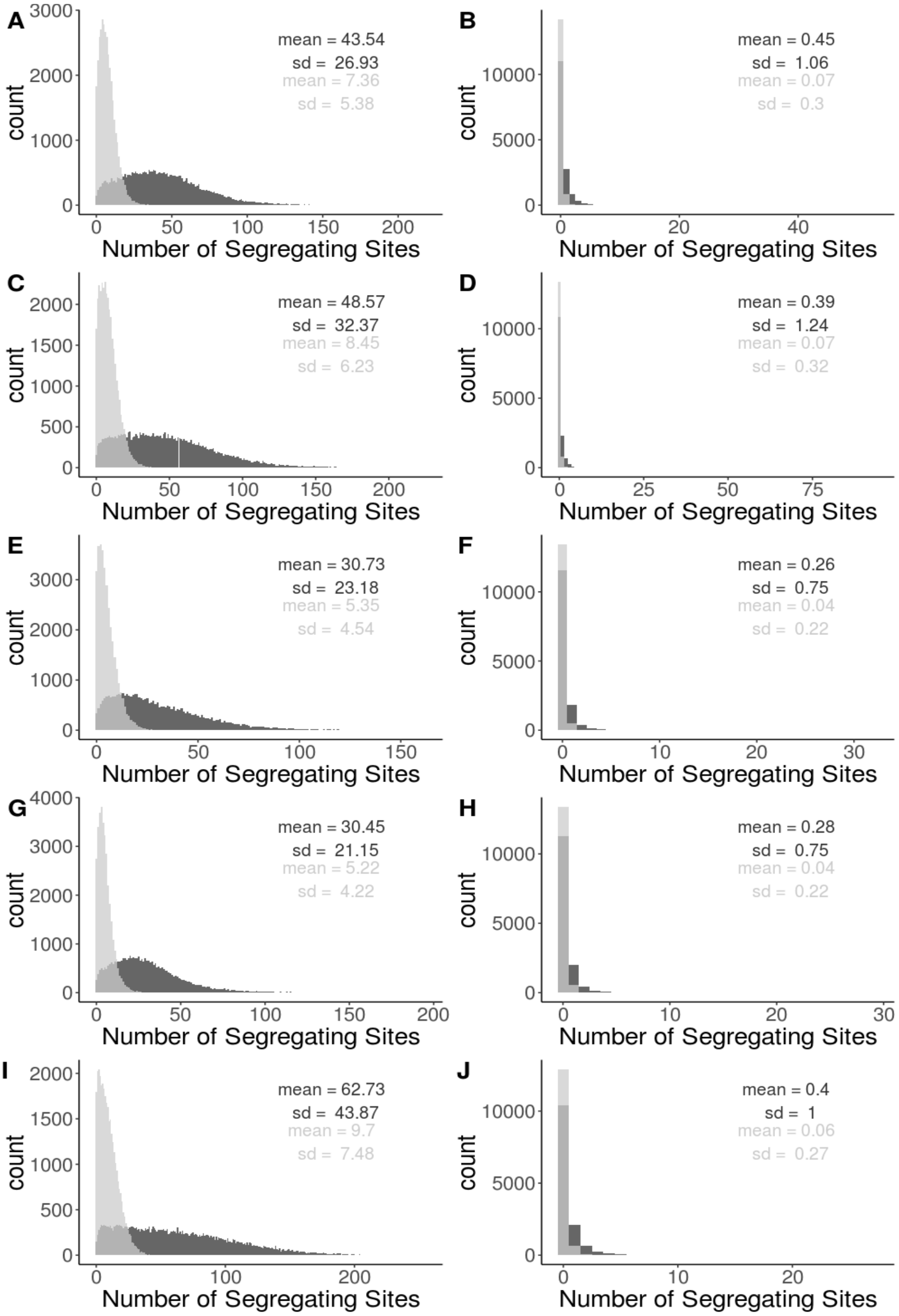


**Supplementary Figure 2**. Distribution of SNP counts in 50KB windows for non-coding sites (left panels) and 0-fold degenerate non-synonymous sites (right panels), respectively, across species. Dark grey for all mutations and light grey for GC-conservative mutations. A-B) humans, C-D) Nigeria-Cameroon chimpanzees, E-F) western chimpanzees, G-H) bonobos and I-J) gorillas.

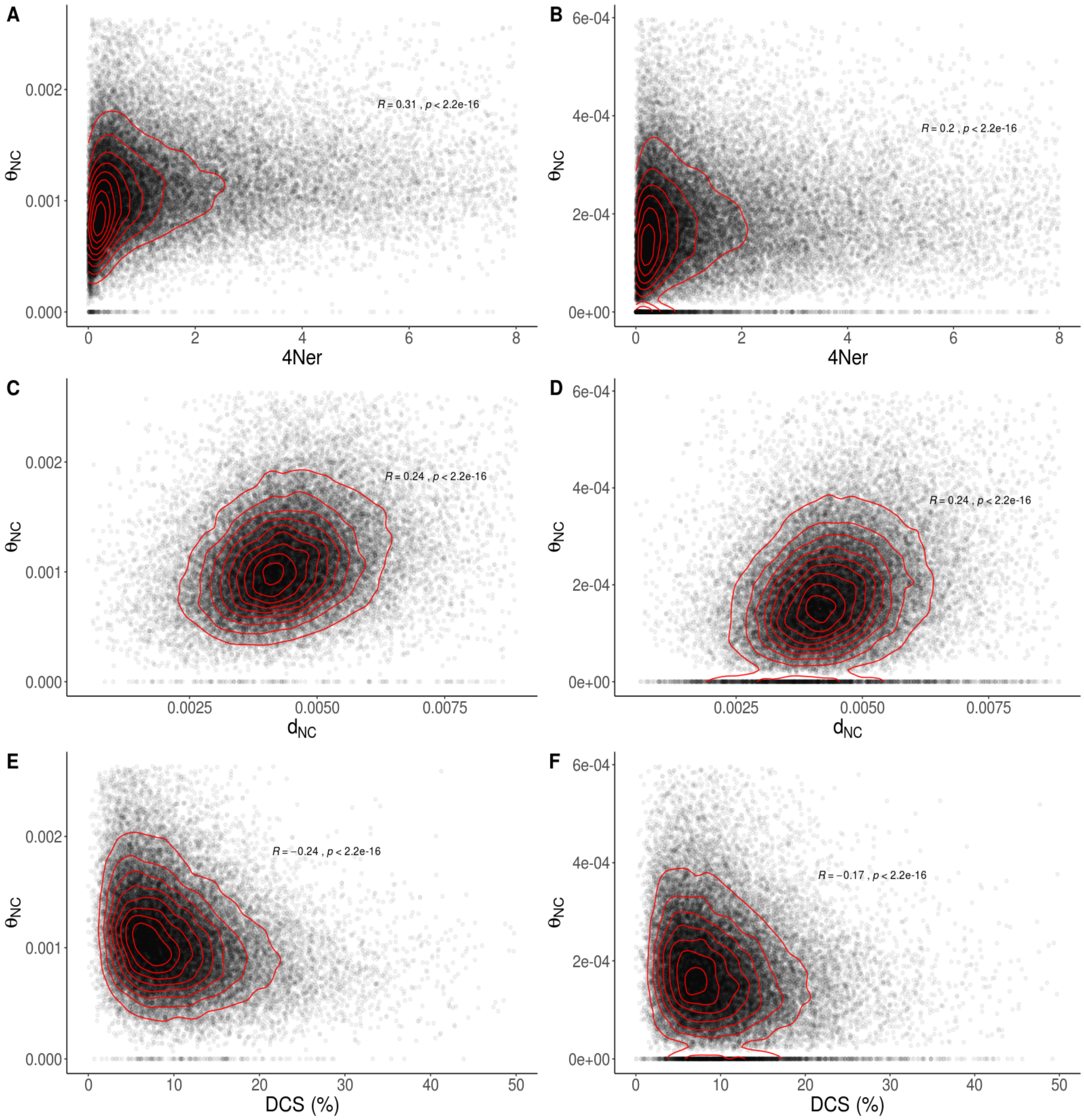


**Supplementary Figure 3**. Relationship between $\theta_{NC}$ and the three genomic features in humans for A, C, E) all mutations and for B, D, F) GC-conservative mutations. Spearman rank correlation coefficients and associated *P*-value. The contour lines in red are added for visualizing the relationship between variables.

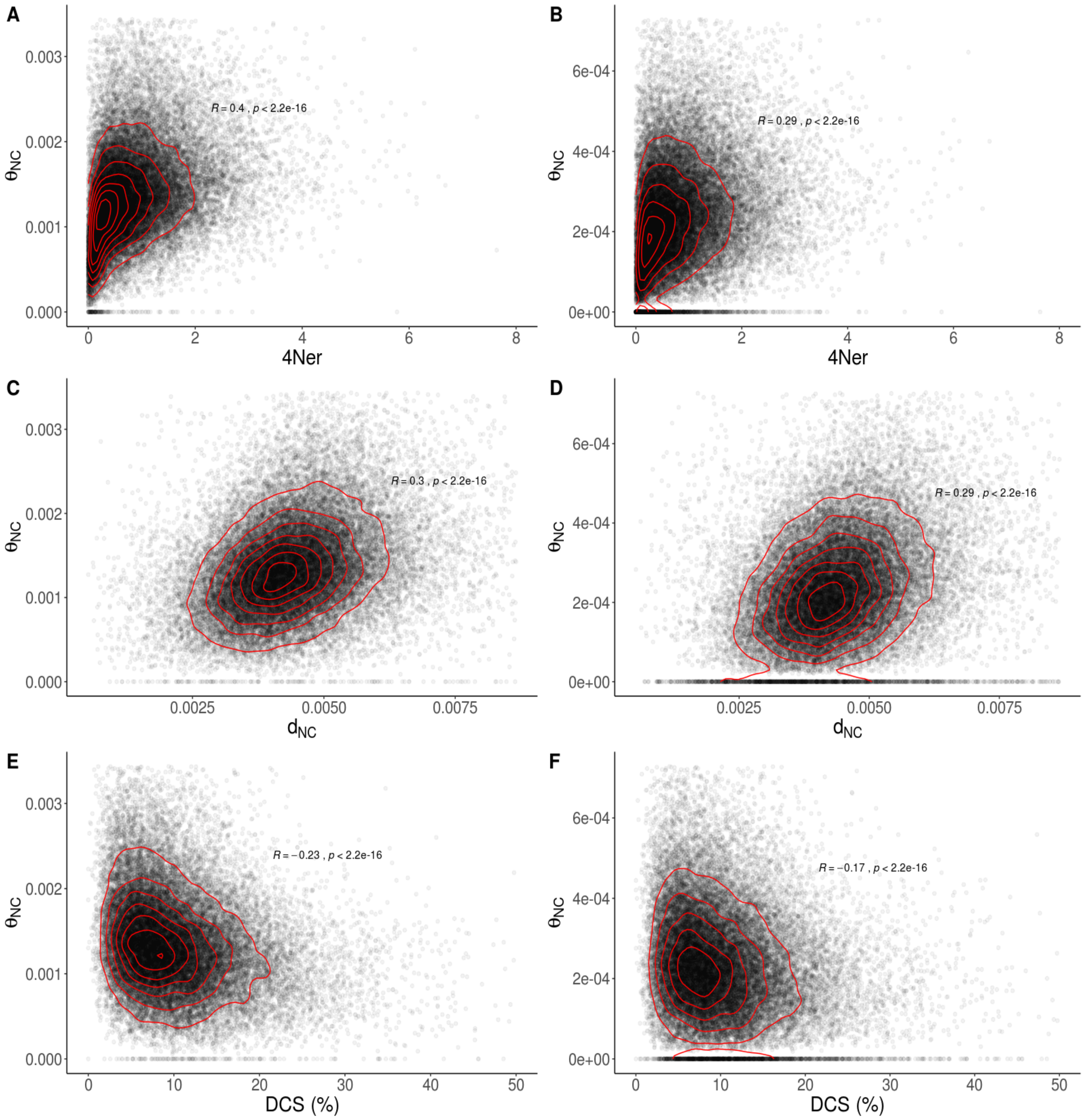


**Supplementary Figure 4**. Relationship between $\theta_{NC}$ and the three genomic features in Nigeria-Cameroon chimpanzees for A, C, E) all mutations and for B, D, F) GC-conservative mutations. Spearman rank correlation coefficients and associated *P*-value. The contour lines in red are added for visualizing the relationship between variables.

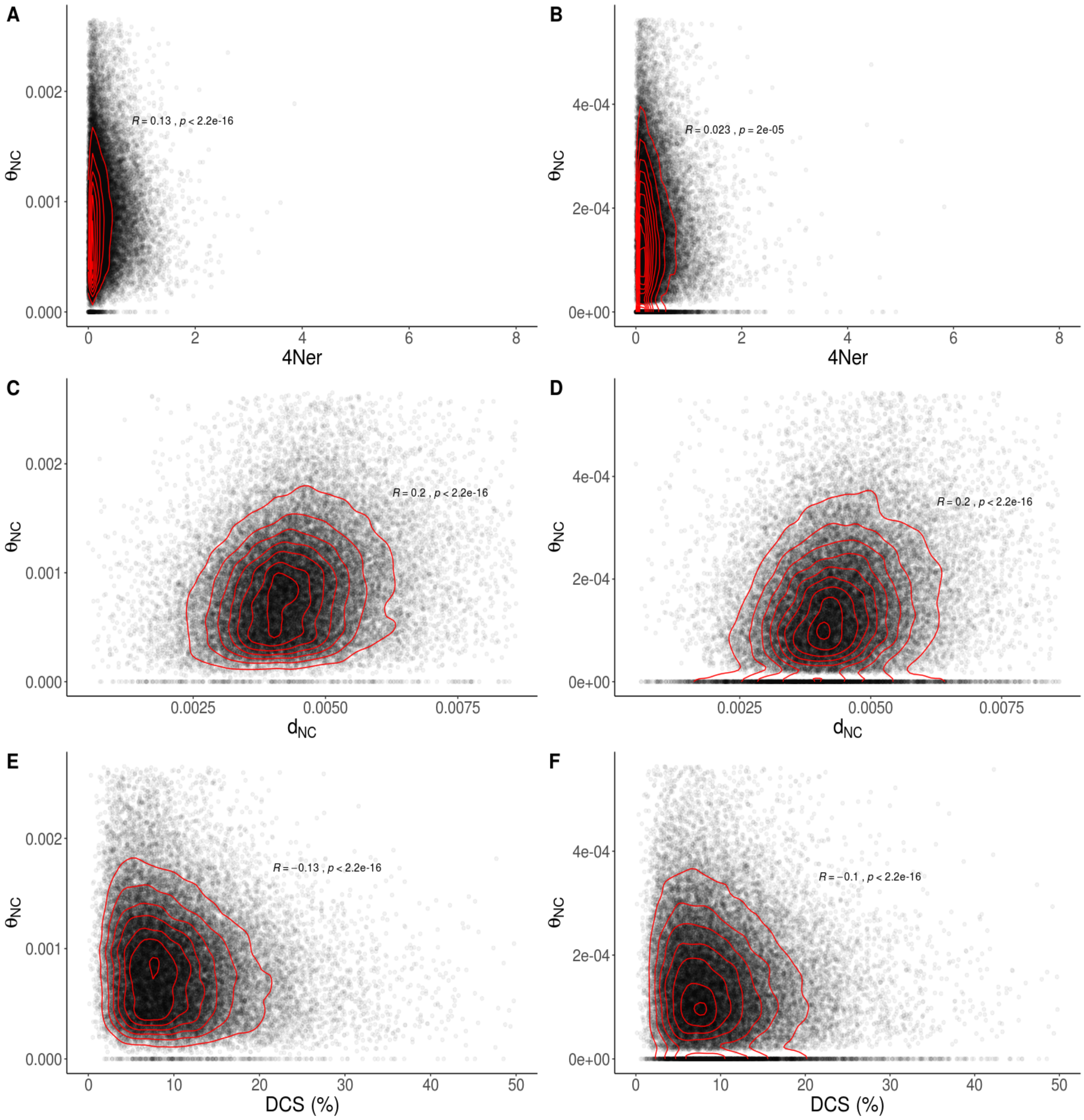


**Supplementary Figure 5**.- Relationship between $\theta_{NC}$ and the three genomic features in western chimpanzees for A, C, E) all mutations and for B, D, F) GC-conservative mutations. Spearman rank correlation coefficients and associated *P*-value. The contour lines in red are added for visualizing the relationship between variables.

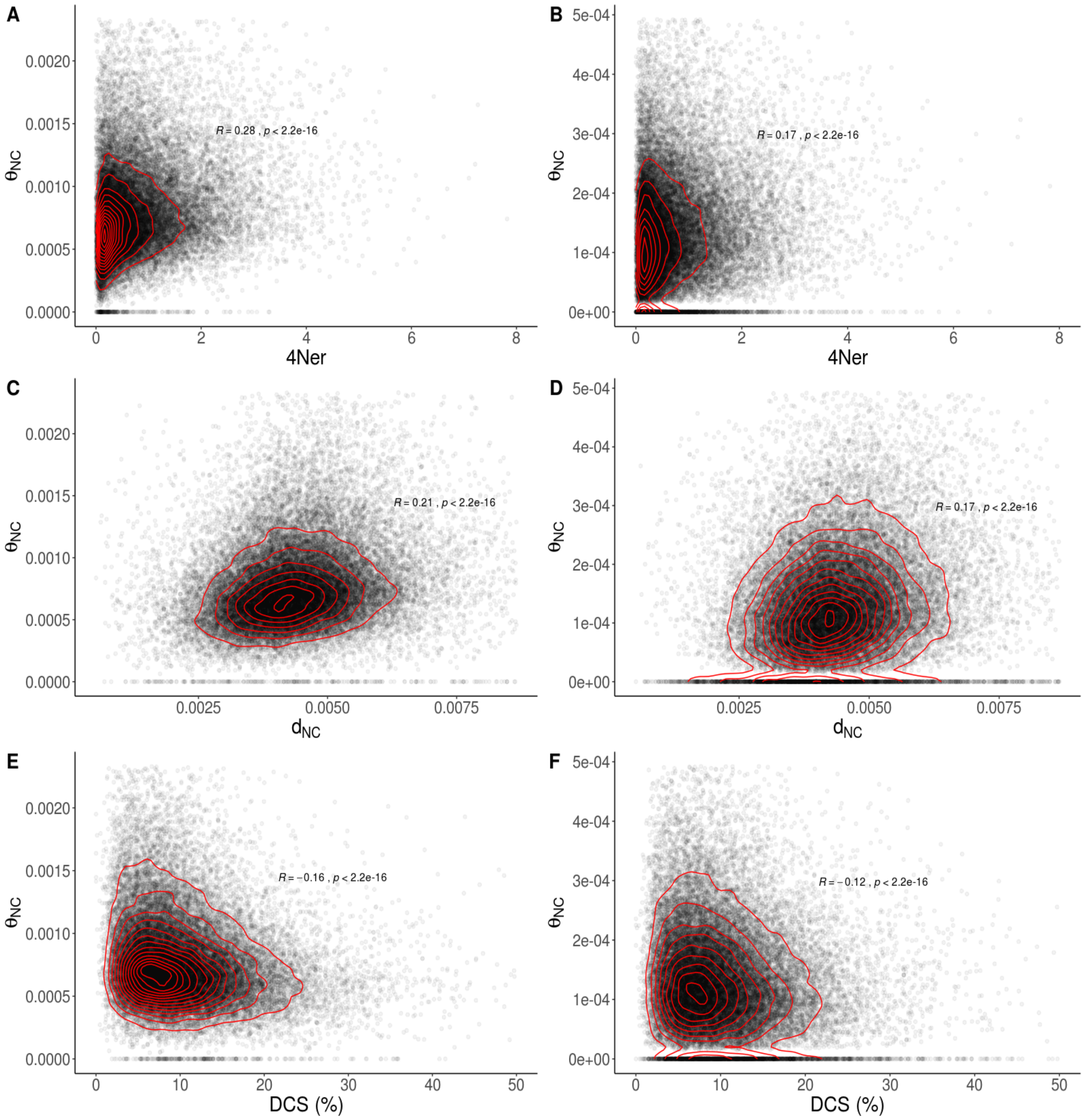


**Supplementary Figure 6**.- Relationship between $\theta_{NC}$ and the three genomic features in bonobos for A, C, E) all mutations and for B, D, F) GC-conservative mutations. Spearman rank correlation coefficients and associated *P*-value. The contour lines in red are added for visualizing the relationship between variables.

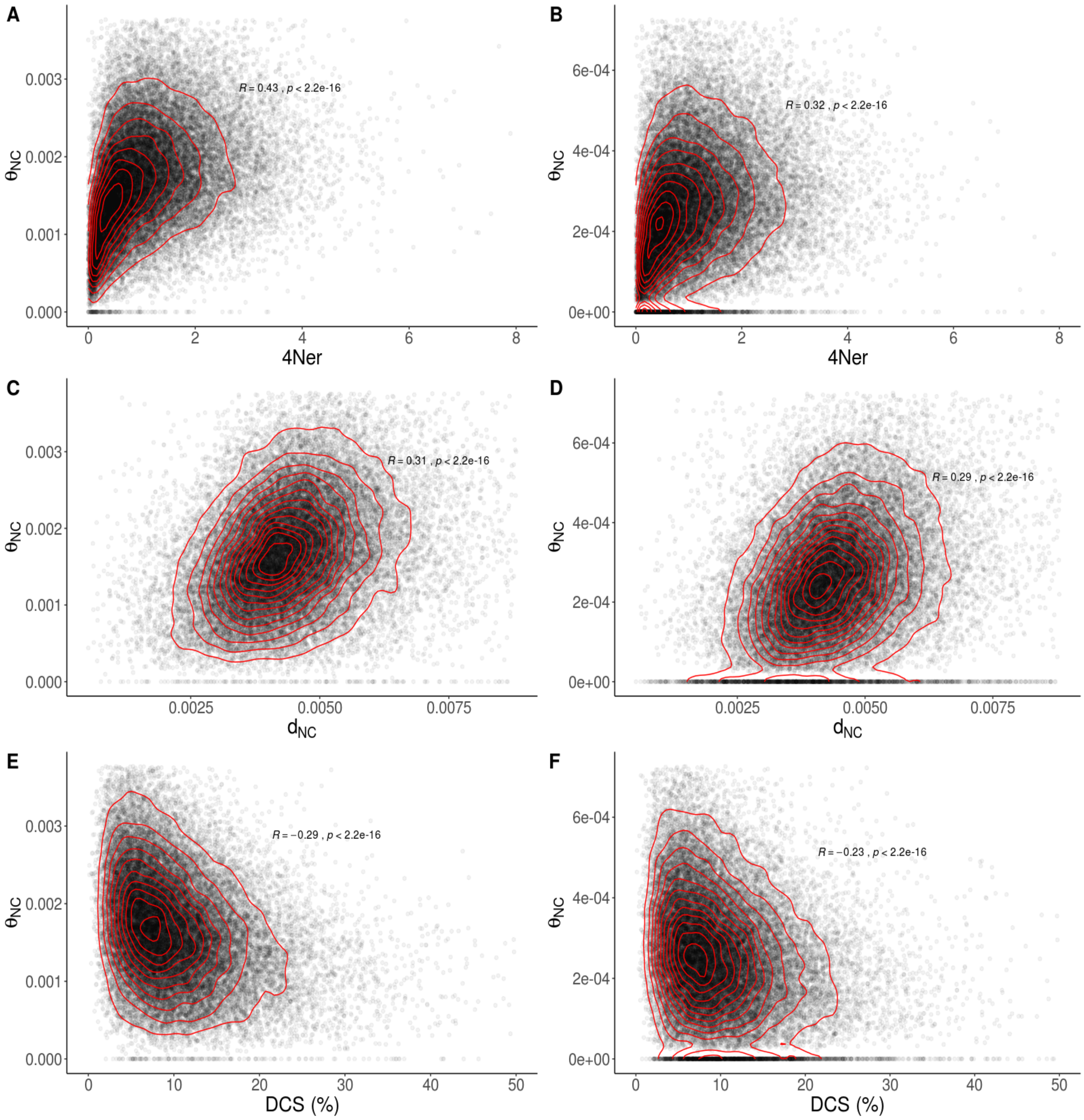


**Supplementary Figure 7**.- Relationship between $\theta_{NC}$ and the three genomic features in gorillas for A, C, E) all mutations and for B, D, F) GC-conservative mutations. Spearman rank correlation coefficients and associated *P*-value. The contour lines in red are added for visualizing the relationship between variables.

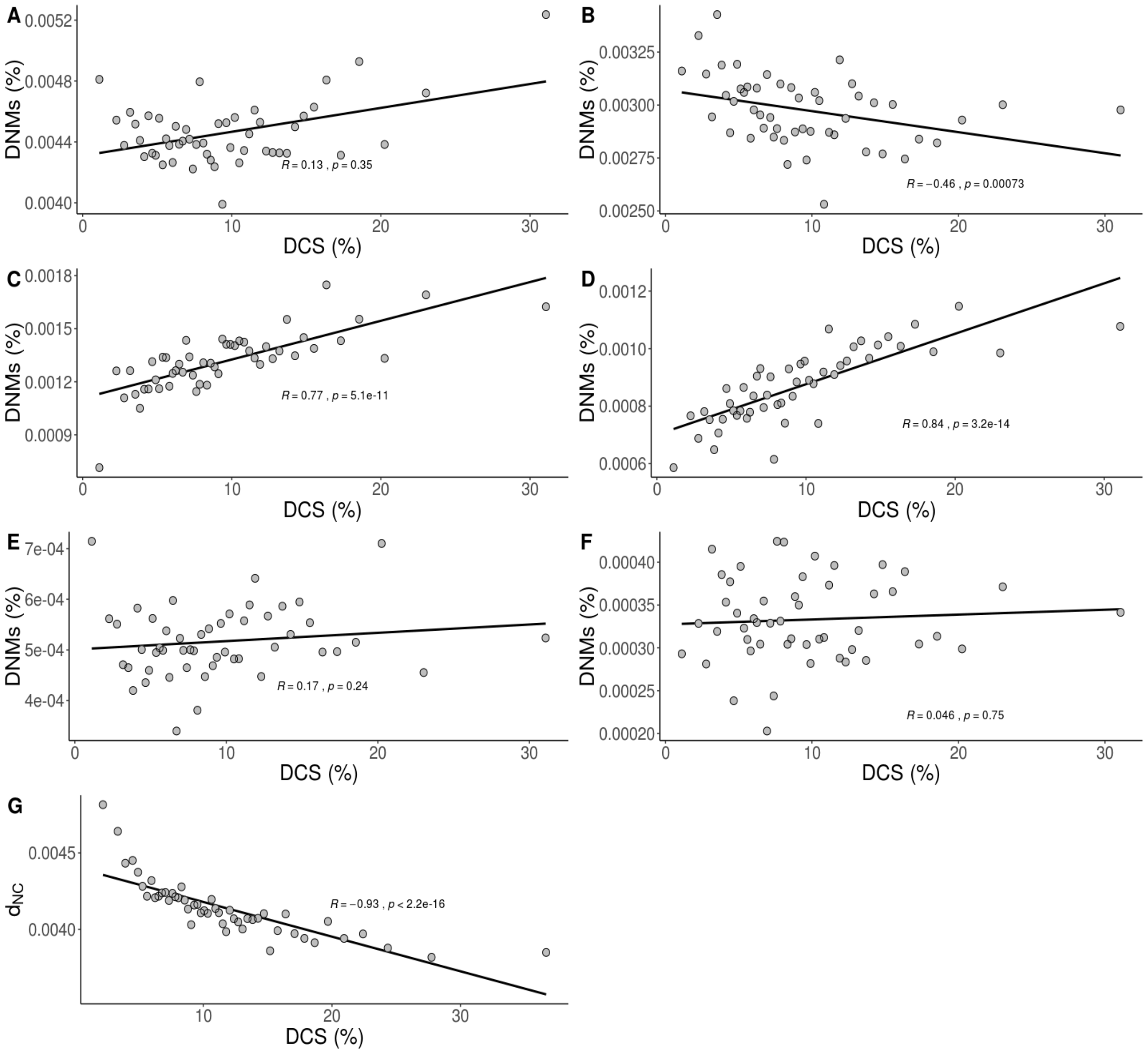


**Supplementary Figure 8.-** Relationship between A, C, E) the rate of putatively neutral DNMs, B, D, F) the rate of putatively selected DNMs and F) $d_{NC}$ and the DCS. DNMs from A-B) Jónsson et al. (2017), C-D) Wong et al. (2016), E-F) Francioli et al. (2015) and G) substitution rates at GC-conservative putatively neutral non-coding sites from this study. The lines are linear regressions along with the Spearman rank correlation coefficients and associated *P*-values. Results after ranking and binning windows by the DCS (into 50 bins).

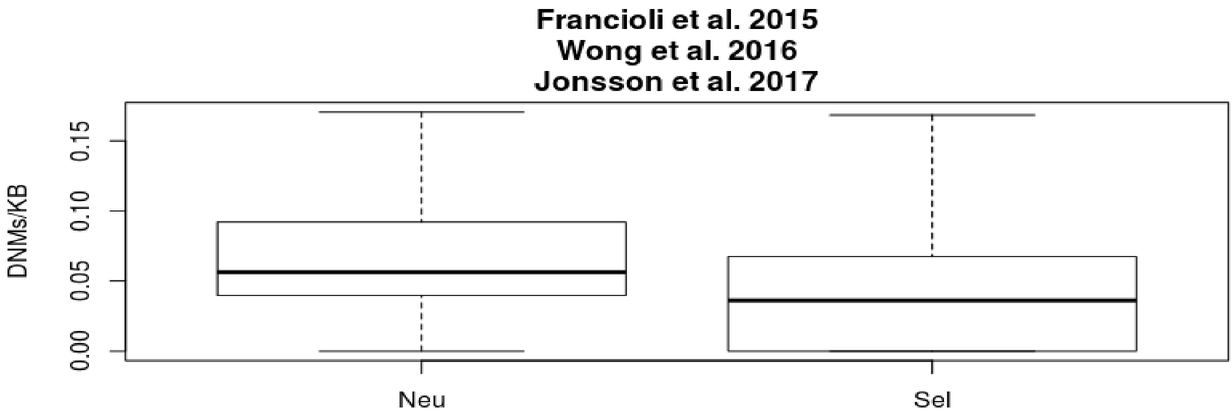


**Supplementary Figure 9.-** Boxplot for the rate of putatively neutral non-CpG DNMs (Neu) and the rate of putatively selected non-CpG DNMs (Sel) across DNM datasets. The outliers have been removed.
