## Supplementary Tables for "Impact of mutation rate and selection at linked sites on DNA variation across the genomes of humans and other homininae"

A)

| | Non-coding | | Non-synonymous (0-fold degenerate) | |
|---|---|---|---|---|
| **Species** | **SNPs** | **L** | **SNPs** | **L** |
| **Human** | 1,529,027 | 527.5 | 6,820 | 7.8 |
| **NC Chimpanzee** | 1,637,818 | 442.0 | 5,616 | 6.4 |
| **W Chimpanzee** | 1,031,001 | 441.8 | 3,591 | 6.3 |
| **Bonobo** | 1,015,810 | 490.2 | 3,845 | 6.4 |
| **Gorilla** | 2,069,332 | 442.3 | 5,536 | 5.5 |

B)

| | Non-coding | | Non-synonymous (0-fold degenerate) | |
|---|---|---|---|---|
| **Species** | **SNPs** | **L** | **SNPs** | **L** |
| **Human** | 258,985 | 528.3 | 1,100 | 7.8 |
| **NC Chimpanzee** | 286,342 | 443.2 | 1,000 | 6.4 |
| **W Chimpanzee** | 180,462 | 442.9 | 596 | 6.2 |
| **Bonobo** | 175,168 | 491.7 | 612 | 6.4 |
| **Gorilla** | 321,993 | 443.9 | 850 | 5.4 |

**Supplementary Table 1.** Total number of analyzed sites (L in MB) and SNPs for A) all mutations and B) GC-conservative mutations.

| *r* \ *P*-Value | Human | NC Chimpanzee | W Chimpanzee | Bonobo | Gorilla | All |
|---|---|---|---|---|---|---|
| **Human** | [redacted] | *** | *** | *** | *** | *** |
| **NC Chimpanzee** | 0.53 | [redacted] | *** | *** | *** | *** |
| **W Chimpanzee** | 0.55 | 0.43 | [redacted] | *** | *** | *** |
| **Bonobo** | 0.65 | 0.45 | 0.46 | [redacted] | *** | *** |
| **Gorilla** | 0.79 | 0.54 | 0.56 | 0.66 | [redacted] | *** |
| **All** | 0.91 | 0.61 | 0.63 | 0.75 | 0.93 | [redacted] |

**Supplementary Table 2**. Spearman correlation coefficients between lineage-specific $d_{NC}$ for all homininae pairs. All represents the substitution rate across the whole tree, this is the proxy of the mutation rate that we use through the study.

| **Species** | **$d_{NC}$** | **DCS** |
|---|---|---|
| **NC Chimpanzee** | 0.15 *** | -0.10 *** |
| **W Chimpanzee** | 0.11 *** | -0.03 *** |
| **Bonobo** | 0.08 *** | -0.06 *** |
| **Gorilla** | 0.15 *** | -0.14*** |

**Supplementary Table 3**. Spearman correlation between RR and $d_{NC}$ and RR and the density of conserved sites (DCS) in non-human homininae.

A)

| Species | RR | $d_{NC}$ | DCS | $R^2$ | $R^2$,L- | $R^2$,L+ |
|---|---|---|---|---|---|---|
| **Human** | 0.0030 *** | 0.0023 *** | -0.0026 *** | 67% | 12% | 63% |
| **NC chimpanzee** | 0.0032 *** | 0.0026 *** | -0.0019 *** | 71% | 14% | 67% |
| **W chimpanzee** | 0.0008 *** | 0.0037 *** | -0.0019 *** | 54% | 7% | 52% |
| **Bonobo** | 0.0044 *** | 0.0025 *** | -0.0019 *** | 68% | 10% | 65% |
| **Gorilla** | 0.0024 *** | 0.0016 *** | -0.0017 *** | 74% | 14% | 70% |

B)

| Species | RR | $d_{NC}$ | DCS | $R^2$ | $R^2$,L- | $R^2$,L+ |
|---|---|---|---|---|---|---|
| **Human** | 0.0988 *** | 0.0561 *** | -0.2817 *** | 27% | 0.64% | 24% |
| **NC chimpanzee** | 0.1047 *** | 0.0605 *** | -0.2460 *** | 27% | 0.62% | 23% |
| **W chimpanzee** | 0.0748 ** | 0.1029 *** | -0.3728 *** | 23% | 0.40% | 21% |
| **Bonobo** | 0.1128 *** | 0.0085 | -0.3703 *** | 24% | 0.47% | 22% |
| **Gorilla** | 0.1034 *** | 0.0660 *** | -0.3714 *** | 25% | 0.16% | 22% |

**S**

**Supplementary Table 4.** The standardised regression coefficients from the full model regressing the number of SNPs against the recombination rate (RR), $d_{NC}$ and density of conserved sites (DCS) along with the $R^2$ for the full model, and a $R^2$ for the model without L and with only L, where L is the number of analyzed sites for A) non-coding SNPs and B) non-synonymous SNPs. * *P*-value<0.05, ** *P*-value<0.01, *** *P*-value<0.001.

A)

| Species | RR | $d_{NC}$ | DCS | $R^2$ | $R^2$,L- | $R^2$,L+ |
|---|---|---|---|---|---|---|
| **Human** | 0.0119 *** | 0.0227 *** | -0.0104 *** | 53% | 11% | 49% |
| **NC chimpanzee** | 0.0142 *** | 0.0212 *** | -0.0076 *** | 60% | 14% | 55% |
| **W chimpanzee** | 0.0014 | 0.0301 *** | -0.0073 *** | 41% | 7% | 39% |
| **Bonobo** | 0.0199 *** | 0.0204 *** | -0.0083 *** | 48% | 8% | 46% |
| **Gorilla** | 0.0120 *** | 0.0152 *** | -0.0089 *** | 65% | 14% | 61% |

B)

| Species | RR | $d_{NC}$ | DCS | $R^2$ | $R^2$,L- | $R^2$,L+ |
|---|---|---|---|---|---|---|
| **Human** | 0.2374 * | 0.1669 | -1.0218 *** | 16% | 0.41% | 14% |
| **NC chimpanzee** | 0.3491 ** | 0.2527 * | -1.0033 *** | 18% | 0.42% | 16% |
| **W chimpanzee** | 0.204 | 0.3047 | -1.4525 *** | 14% | 0.26% | 13% |
| **Bonobo** | 0.3335 * | 0.1897 | -1.3009 *** | 14% | 0.24% | 13% |
| **Gorilla** | 0.2977 * | 0.1518 | -1.6346 *** | 14% | 0.07% | 12% |

**S**

**Supplementary Table 5.** The standardised regression coefficients from the full model regressing the number of GC-conservative SNPs against the recombination rate (RR), $d_{NC}$ and the density of conserved sites (DCS) along with the $R^2$ for the full model, and a $R^2$ for the model without L and with only L, where L is the number of analyzed sites for A) non-coding SNPs and B) non-synonymous SNPs. * *P*-value<0.05, ** *P*-value<0.01, *** *P*-value<0.001.

**A)**

| **Species** | **RR** | **$d_{NC}$** | **DCS** |
|---|---|---|---|
| **Human** | 0.03 (-0.38, 0.46) | -0.13 (-0.55, 0.29) | 0.10 (-0.37, 0.57) |
| **NC Chimpanzee** | -0.15 (-0.75, 0.41) | -0.05 (-0.53, 0.42) | 0.00 (-0.38, 0.40) |
| **W Chimpanzee** | -0.27 (-0.54, -0.02) | -0.07 (-0.34, 0.23) | 0.02 (-0.27, 0.33) |
| **Bonobo** | -0.21 (-0.58, 0.15) | -0.18 (-0.53, 0.19) | -0.20 (-0.46, 0.05) |
| **Gorilla** | 0.02 (-0.52, 0.55) | 0.38 (-0.05, 0.82) | *0.58 (0.13, 0.98)* |

**B)**

| **Species** | **RR** | **$d_{NC}$** | **DCS** |
|---|---|---|---|
| **Human** | -0.31 (-0.64, 0.05) | -0.05 (-0.34, 0.23) | -0.10 (-0.41, 0.23) |
| **NC Chimpanzee** | -0.07 (-0.53, 0.40) | -0.10 (-0.48, 0.28) | -0.04 (-0.38, 0.30) |
| **W Chimpanzee** | 0.06 (-0.22, 0.34) | -0.10 (-0.30, 0.09) | 0.03 (-0.25, 0.31) |
| **Bonobo** | -0.16 (-0.45, 0.12) | -0.07 (-0.40, 0.27) | 0.03 (-0.24, 0.29) |
| **Gorilla** | -0.14 (-0.55, 0.26) | -0.10 (-0.50, 0.27) | -0.25 (-0.59, 0.10) |

**Supplementary Table 6.** The standardised regression coefficients of a multiple linear regression of log ($\theta_N/\theta_{NC}$) versus RR, $d_{NC}$ and the the density of conserved sites (DCS) A) for all SNPs and B) for GC-conservative SNPs. To assess the robustness of our standardised regression coefficients in the presence of multicollinearity we bootstrapped the genome by window one-thousand times. Given in parentheses are the 95% confidence intervals by bootstrapping.
